# An adapted method for Cas9-mediated editing reveals the species-specific role of β-glucoside utilization driving competition between *Klebsiella* species

**DOI:** 10.1101/2023.09.27.559776

**Authors:** Éva d. H. Almási, Nele Knischewski, Lisa Osbelt, Uthayakumar Muthukumarasamy, Youssef El Mouali, Elena Vialetto, Chase L. Beisel, Till Strowig

## Abstract

Cas9-based gene editing tools have revolutionized genetics, enabling the fast and precise manipulation of diverse bacterial species. However, widely applicable genetic tools for non-model gut bacteria are unavailable. Here, we present a two-plasmid Cas9-based system designed for gene deletion and knock-in complementation in three members of the *Klebsiella oxytoca* species complex (KoSC), which we applied to study the genetic factors underlying the role of these bacteria in competition against *Klebsiella pneumoniae*. The system allowed efficient and precise editing via enhanced lambda Red expression and functionally-validated complementation with the use of universal ‘bookmark’ targets in *K. oxytoca, Klebsiella michiganensis*, and *Klebsiella grimontii*. We revealed that the carbohydrate permease CasA is critical in *ex vivo* assays for *K. pneumoniae* inhibition by *K. oxytoca* but is neither sufficient nor required for *K. michiganensis* and *K. grimontii*. Thus, the adaptation of state-of-the-art genetic tools to KoSC allows the identification of species-specific functions in microbial competition.

**\* Importance:** Cas9-based gene editing tools have revolutionized bacterial genetics, yet, their application to non-model gut bacteria is frequently hampered by various limitations. We utilized a two-plasmid Cas9-based system designed for gene deletion in *Klebsiella pneumoniae* and demonstrate after optimization its utility for gene editing in three members of the *Klebsiella oxytoca* species complex (KoSC) namely *K. oxytoca, K. michiganensis* and *K. grimontii*. We then adapted a recently developed protocol for functional complementation based on universal ‘bookmark’ targets applicable to all tested species. In summary, species specific adaptation of state-of-the-art genetic tools allow efficient gene deletion and complementation in types strains as well as natural isolates of KoSC members to study microbial interactions.

## Introduction

An increasing number of multidrug-resistant (MDR) pathogens pose a major health threat worldwide. *Klebsiella pneumoniae*, a common MDR opportunistic pathogen from the family Enterobacteriaceae^1^, is estimated to have a prevalence of 32.8% among nosocomial infection isolates worldwide^2^. As the discovery of novel antimicrobials has proven to be both increasingly difficult and slow^3^, many recent studies have explored alternatives to classic antibiotics, specifically, how the microbiome can be utilized to prevent infections^4–8^. It is now widely recognized that the microbiome contributes to colonization resistance (CR) against disease-causing pathogens^9,10^ through various mechanisms. For instance, commensals can protect the host by utilizing growth-limiting carbon sources^5–7^, siderophores^11^, and oxygen consumption^8^ or by producing bioactive molecules such as secondary bile acids^4^, short-chain fatty acids^12^ and growth inhibitory toxins^13,14^. Diverse human-isolated commensal bacteria have also been described to enhance CR, either by preventing pathogen outgrowth or facilitating pathogen clearance. Such species include enterobacteria, such as *E. coli*^7,8^ and members of the *Klebsiella oxytoca* species complex (KoSC)^5,6^. For instance, *K. oxytoca* was shown to enhance colonization resistance against *K. pneumoniae* in ampicillin-treated mice via competition for β-glucosides, which was dependent on the β-glucoside transporter, CasA^6^. As in many cases, the mechanisms through which pathogen elimination occurs vary depending on the interacting microbes, and additional functional studies are necessary to gain insights into which properties are critical for optimal exploration as future therapeutics.

Genetic tools in bacteria have been decisive in the mechanistic dissection of bacterial physiology and microbial interactions. For gut commensals, such efforts have vastly focused on model organisms such as *E. coli*^15^ or *B. thetaiotaomicron*^16^, whereas for many other commensal bacteria, these tools have been lacking. Notably, the speed and efficiency of genetic engineering has been greatly improved by the development of CRISPR technologies in the past decade^17,18^. For bacterial gene editing, CRISPR-Cas9 systems are frequently used together with phage-derived recombination systems^19^, typically with lambda Red^20–22^. Similar to traditional recombineering approaches, a DNA template is provided to the recipient cells to repair the chromosomal lesion via homologous recombination facilitated by a recombination system. However, while recombineering relies on the incorporation of an antibiotic resistance marker into the chromosome for selection of edited cells^23^, in the case of CRISPR/Cas9-based tools, the double-stranded break (DSB) by Cas9 counter-selects against unedited cells, as DBS likely leads to cell death in bacteria^19^. Despite many significant iterative technological advances for model organisms, it has been challenging to adopt them in less-studied species and non-laboratory strains of gut bacteria.^24^

Here, we describe the adaptation and enhancement of a CRISPR-Cas9 tool coupled with lambda Red recombination in both human isolates and type strains of three species of the KoSC: *K. oxytoca, K. michiganensis* and *K. grimontii*. Characterizing the role of KoSC in the gut has attracted growing interest, as it has been reported to enhance colonization resistance^5,6^ as well as cause nosocomial infections^25,26^, but so far genetic tools have only been reported for gene knockout in *K. oxytoca*^6,23,25^. The optimized protocol allows full-length gene deletion and knock-in complementation without the long-term maintenance of antibiotic-resistance genes, which enables plasmid-free *ex vivo* and *in vivo* studies with engineered strains. Furthermore, as a proof-of-concept, we show how competition for β-glucosides as a driving force behind the decolonization of *K. pneumoniae* is species-specific to *K. oxytoca*, rather than being a general principle in the KoSC.

## Results

### Strong induction of homologous recombination is required for high-efficiency gene deletion

Our adaptation and optimization strategy was built on a two-plasmid CRISPR-Cas9 based system originally developed in *K. pneumoniae*^21^, which we recently utilized to target genes in the *K. oxytoca* strain MK01^6^. Briefly, the endonuclease Cas9 was encoded on the same plasmid as the phage-derived lambda Red recombination system, which is under the transcriptional control of the L-arabinose-inducible *P*_*araB*_ promoter (Figure 1a). The second plasmid encodes the Cas9 handle of a single-guide RNA (sgRNA) and *sacB*^27^, a negative selection that can be used for plasmid curing (Figure 1a). We modified this plasmid to encode a GFP dropout sequence upstream of the Cas9 handle to allow rapid selection of correctly ligated plasmids containing the guide of interest. The third component is a linear double-stranded DNA repair template assembled from 500-bp homology arms carrying the desired genetic edit. Gene editing was achieved by iterative transformation of the first Cas9-encoding plasmid (pCas-apr), followed by the sgRNA-encoding plasmid (pSG-spe), and the repair template.

**Figure 1:**
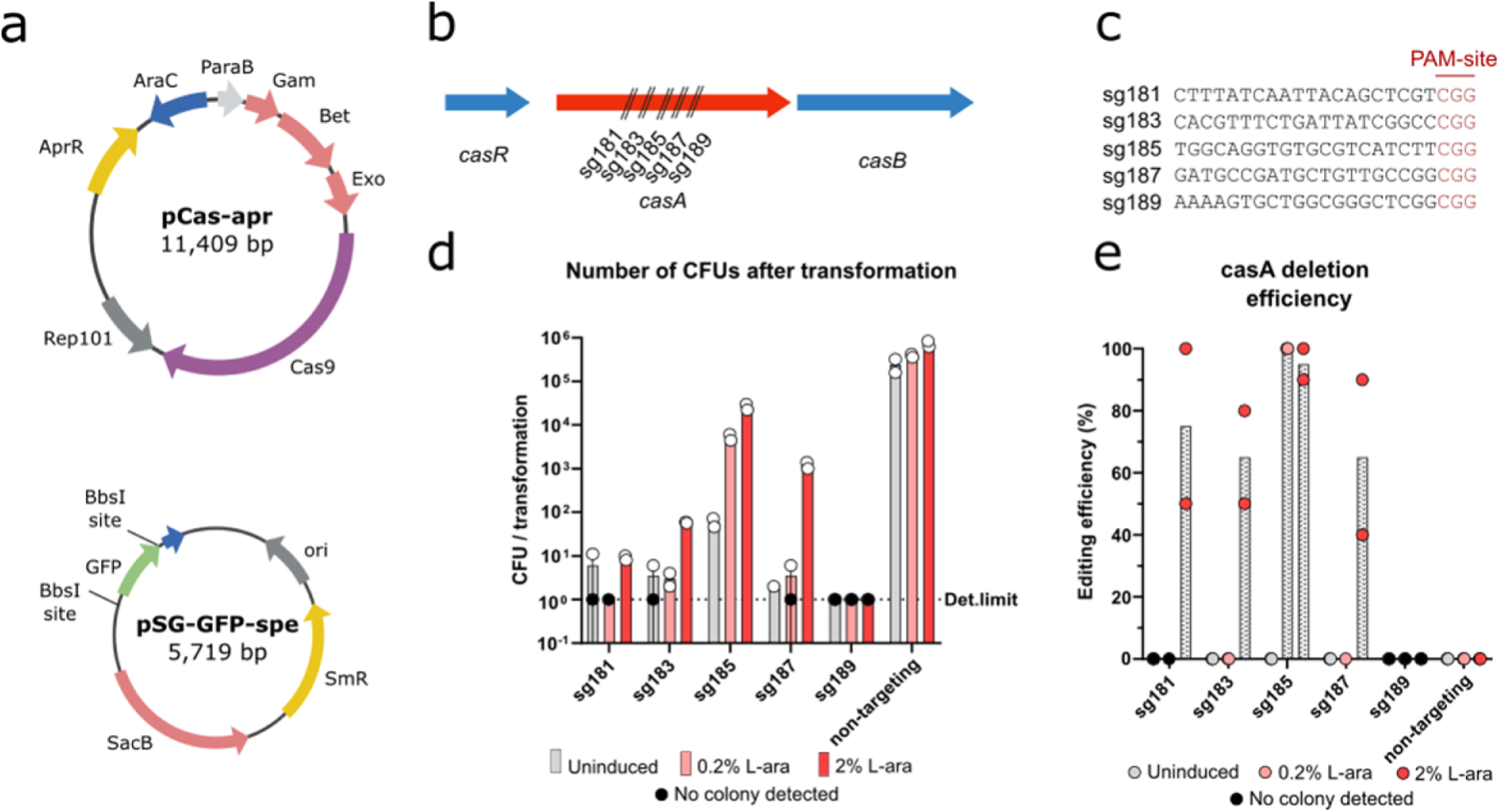
Two-plasmid CRISPR-Cas9 system allows highly efficient deletion of *casA*. **(a)** Plasmid maps of pCas-apr and pSG-GFP-spe. **(b)** Schematic and **(c)** sequences of spacers designed for *casA* deletion **(d)** Number of colonies under different lambda Red induction conditions. Under uninduced conditions no repair template was used in order to show killing efficiency. 0.2% and 2% L-arabinose-induced cells were provided a repair template. Colony counts were compared to transformation of pSG-spe expressing a non-targeting gRNA. All transformations were performed in biological duplicates using separately prepared batches of competent cells. Each dot represents CFUs from one biological replicate **(e)** Deletion efficiencies of different sgRNAs quantified by colony PCR reactions. Individual dots represent average of 10 randomly selected colonies screened from targeting samples and 3 randomly selected colonies screened from non-targeting samples.

To adapt and optimize the protocol, we initially compared 5 *casA*-targeting sgRNAs in their ability to facilitate successful full-length gene deletion in *K. oxytoca* MK01 (Figure 1b-c). First, the targeting capacity of the sgRNAs was assessed by transformation of plasmid pSG-spe into *K. oxytoca* MK01 in the absence of a repair template and without induction of the lambda Red machinery. Under these conditions, CRISPR-Cas9 is toxic to cells owing to its DBS activity. Compared to the non-targeting control, we observed a 10^4^-10^5^-fold reduction in CFUs in four out of five sgRNAs and no detectable colonies with sg189 (Figure 1d), showing that the sgRNAs led to efficient killing in the absence of a repair template with a small number of surviving colonies (Figure 1e), even though both plasmids were present in the cell at the same time.

Next, to promote homologous recombination-driven gene deletion of *casA*, we induced expression of the lambda Red system by adding 0.2% L-arabinose, as reported for *K. pneumoniae*^21^. Surprisingly, we observed an increase in CFU compared to the uninduced conditions for only one of the five sgRNAs, sg185 (approximately 100-fold increase, from 59 to 5250 CFUs) (Figure 1d). The deletion efficiency of sg185 was increased from 0 to 100% as evidenced by colony PCR (10/10 tested colonies), but four out of five sgRNAs led to no successful gene deletion of *casA* (Figure 1e). As the survival of cells in most cases depends on successful recombination, we hypothesized that stronger induction of lambda Red gene expression may lead to better editing with other sgRNAs. Indeed, increasing the L-arabinose concentration ten-fold increased the CFU count from 9 to >10^4^ for three of the four remaining sgRNAs (Figure 1d). More importantly, for these sgRNAs a 65-95% deletion efficiency of *casA* was observed (Figure 1e). Collectively, these results show that while sgRNA performance can vary in terms of both surviving CFUs and editing efficiency, sufficient expression of the lambda Red system leads to robust editing with four out of five sgRNAs.

### Knock-in complementation results in markerless genotype restoration

To adapt the editing protocol to gene complementation, the same general principles applied within the gene deletion pipeline were initially followed, with specific changes that preclude targeting of the repair template by Cas9. In brief, sgRNAs were designed to target either the upstream adjacent gene *casR* or the downstream adjacent gene *casB*, and a repair template containing the deleted gene flanked by homology arms was assembled (Figure 2a). To avoid targeting and subsequent cleavage of the repair template by Cas9, the target and PAM in the repair template were mutated during homology arm amplification using site-directed mutagenesis. After transformation of the repair template and pSG-spe with four different assembled sgRNAs targeting either *casR* or *casB* into 2% L-arabinose-induced Δ*casA* cells, we observed 10^3^-10^5^-fold CFU reduction compared to the non-targeting control with varying numbers of surviving colonies (n= 0-596 CFU/sgRNA), demonstrating once again the different targeting efficiencies of sgRNAs (Figure 2c). In contrast to the gene deletion pipeline, markedly lower recombination efficiency was observed despite induction with 2% arabinose. Specifically, only one out of four sgRNAs showed successful integration of the repair template, with 10% complementation, in contrast to the 65-95% deletion efficiency (Figures 1e, 2d). This complementation approach, although at a lower frequency, allowed the generation of a marker-less complementation strain, for example, *K. oxytoca* MK01 ∆*casA::casA*.

**Figure 2:**
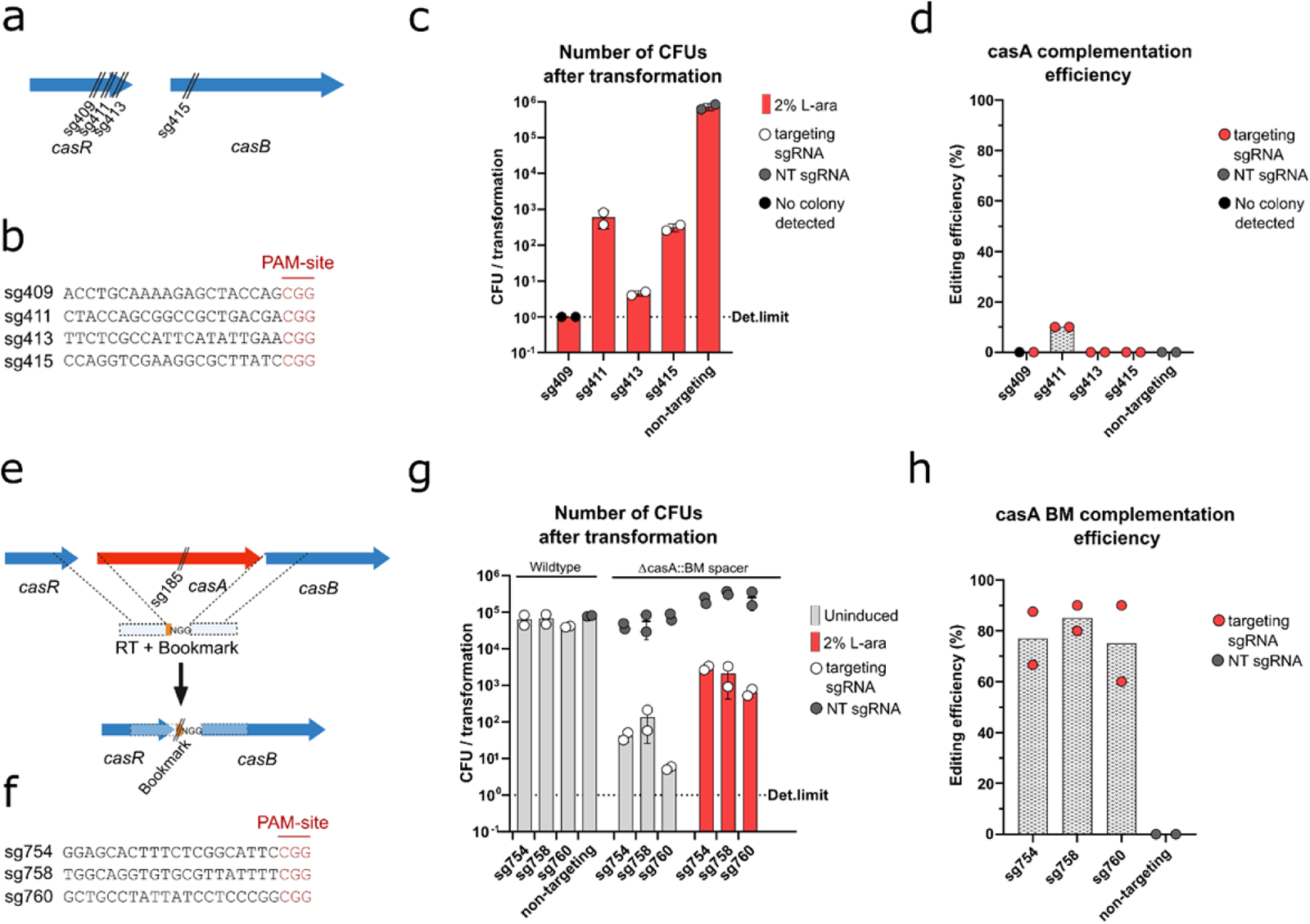
Two knock-in complementation approaches lead to successful genetic complementation of *casA*. **(a, e)** Schematic and **(b, f)** sequences of spacers designed for *casA* complementation and bookmark (BM) complementation, respectively, in *K. oxytoca* MK01. **(c, g)** Number of colonies after transformation of *K. oxytoca* MK01 cells for complementation compared to a non-targeting control. All transformations were performed in biological duplicates using separately prepared batches of competent cells, which were all induced with 2% L-arabinose. Each dot represents CFUs from one biological replicate. **(d, h)** Complementation efficiencies of different sgRNAs. Individual dots represent average of 10 randomly selected colonies screened from targeting samples and 3 randomly selected colonies screened from non-targeting samples.

To overcome the low and target-dependent complementation efficiency observed before, we adapted an alternative complementation method^28^ that is based on the insertion of a 24-bp ‘bookmark’ sequence as part of the repair template during gene deletion, which can then be utilized as a unique sgRNA target with known efficiency during complementation at any genomic locus (Figure 2e). First, we designed four bookmark targets with sequences that were not present in *K. oxytoca* MK01 (Figure 2f) and tested them in uninduced *K. oxytoca* WT cells. No CFU reduction was observed compared to the non-targeting control, suggesting that the bookmark sgRNAs did not cause DBS-triggered cell death in WT cells (Figure 2g). Next, we transformed uninduced Δ*casA::BM* cells with pSG-spe and bookmark sgRNAs to validate targeting after bookmark insertion. Compared to the non-targeting controls, each bookmark sgRNAs showed 10^3^-10^4^-fold reduction in CFUs, confirming successful bookmark insertion into the chromosome and targeting by Cas9 (Figure 2g). Strikingly, following co-transformation of bookmark-ligated pSG-spe plasmids with a repair template containing *casA* and homology arms, we achieved 60-90% complementation efficiency with three out of four tested sgRNAs (Figure 2h). Notably, the bookmark-based complementation approach can be readily used in other genomic loci without prior target-gene specific design. Overall, iterative improvements to the gene editing strategy allowed the highly efficient knock-in of a 2.8 kb insert including *casA* in *K. oxytoca* MK01. This enabled marker-less genomic complementation without the need for a selection marker for plasmid-based complementation.

### CRISPR-Cas9 mediated gene deletion and complementation is applicable across the *Klebsiella oxytoca* species complex (KoSC)

Next, we tested the efficacy of the deletion pipeline in additional strains belonging to the KoSC. We included the type strains for all three KoSC species (*K. oxytoca* DSM5175^T^, *K. michiganensis* DSM25444^T^ and *K. grimontii* DSM105630^T^) and one commensal strain for each KoSC species (*K. michiganensis* LK158 and *K. grimontii* LK33) that was recently isolated from the stool of healthy human donors (Figure 3a). For these five strains, sgRNAs sg185 and sg758 with the highest efficiency for *casA* deletion and subsequent complementation in *K. oxytoca* MK01, respectively, were evaluated. For sg185, alignment of the targets and flanking PAM sequences to genomic sequences revealed 100% sequence identity between the *K. oxytoca* and *K. michiganensis* strains but two mismatches between both *K. grimontii* strains and sg185 (Figure 3f). Notably, Cas9 has been reported to retain functionality despite mismatches between the sgRNA guide and genomic target^29^, and these mismatches were not anticipated to preclude successful targeting in *K. grimontii*. A non-targeting guide RNA encoding pSG-spe was included as a control for transformation efficiency. Notably, the transformation efficiency varied for the five strains between 385 and 5×10^6^ CFU/transformation (Figure 3b). Notably, the transformation efficiency appeared to be strain-specific rather than species-specific (Figure 3b). Colony PCR confirmed successful deletion of *casA* in four out of five tested species ranging from to 10-80% in deletion efficiency, demonstrating the robust applicability of our gene deletion method in KoSC in both type strains and human isolates (Figure 3c).

**Figure 3:**
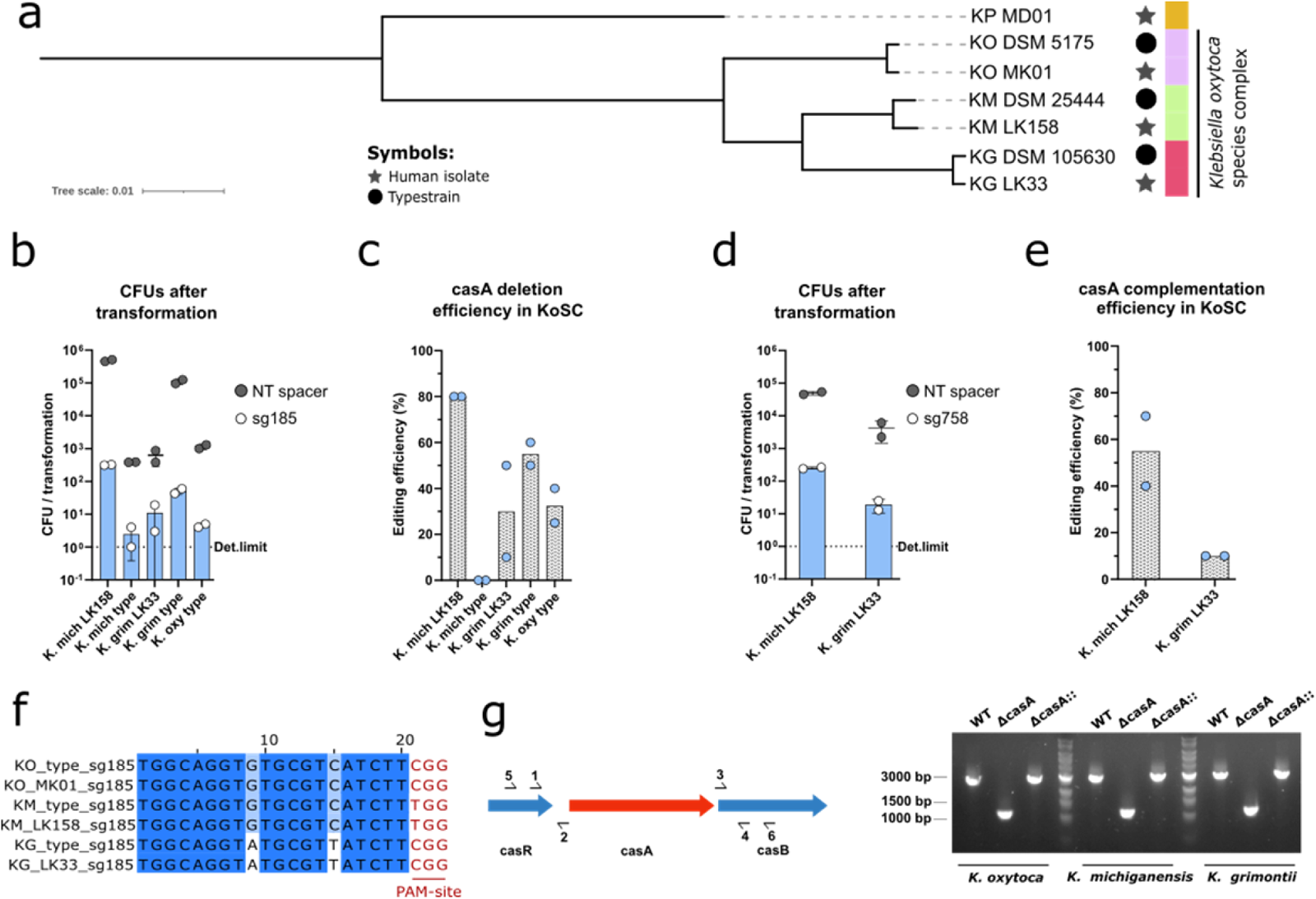
Deletion and complementation protocol extends to other KoSC species. **(a)** Phylogenomic tree representing the evolutionary relations in the KoSC. Symbols indicate the source of the strains. Tree was constructed based on the core genes in the indicated 7 genomes (242 genes). **(b, d)** Number of colonies after transformation of **(b)** pSG-sp185 for *casA* deletion and **(d)** pSG-sp758 for *casA* complementation compared to a non-targeting control. All transformations were performed in biological duplicates using separately prepared batches of competent cells induced with 2% L-arabinose. Each dot represents CFUs from one biological replicate. **(c, e)** Editing efficiencies of **(c)** sp185 and **(e)** for sp758. Individual dots represent average of 10 randomly selected colonies screened from targeting samples. **(f)** Sequence alignment sp185 targeting *casA* for gene deletion in 6 KoSC strains. **(g)** Left: schematic of primers used in casA deletion repair template assembly (1-4) and PCR confirmation of mutant and complemented genotypes (5-6). Right: Gel image of PCR amplicons of WT, *ΔcasA* and *ΔcasA::casA* strains in three KoSC members, expected band size for *ΔcasA* amplicon is ~1100 bp and ~3000 bp for WT and *ΔcasA::casA* amplicons.

Next, we sought to complement the isolated strains of *K. michiganensis* and *K. grimontii*. After selecting sg758 for bookmark complementation because of its robust complementation efficiency in *K. oxytoca* MK01, we genetically deleted *casA* while inserting the sg758 target sequence into the chromosomes of *K. michiganensis* LK158 and *K. grimontii* LK33 to obtain Δ*casA::sg758* cells. Subsequent transformation of pSG-sg758 into Δ*casA::sg758* cells resulted in a 100-fold CFU reduction compared with the non-targeting control, indicating successful targeting (Figure 3d). Furthermore, 50% and 10% complementation efficiency were achieved in *K. michiganensis* LK158 and *K. grimontii* LK33, respectively (Figure 3e), demonstrating that the ‘bookmark’ complementation method is successfully transferrable to different strains. Of note, the editing efficiency of the same sgRNA is highly variable across different strains, with those yielding fewer transformants, generally leading to poorer editing (Figure 3c, e).

### CasA contributes to competition against *K. pneumoniae* only in *K. oxytoca*, not across KoSC

*casA* is part of the *casRAB* operon, which has been previously identified to be required for the utilization of β-glucosides, such as salicin, arbutin, and cellobiose, in *K. oxytoca*^30^. In *K. oxytoca* MK01, we identified *casA* as essential for the utilization of salicin and arbutin, but not cellobiose, and this gene contributes to nutrient competition with multidrug-resistant *K. pneumoniae* strains *ex vivo* and *in vivo*^6^. Since *casA* is present in the KoSC strains tested, we characterized how the WT, Δ*casA* and Δ*casA*::*casA* strains of all three selected species utilize these carbohydrates. In the minimal medium, the utilization of salicin and arbutin was *casA*-dependent in all three species, whereas cellobiose utilization was *casA*-independent (Figure 4a). For all three species five different Δ*casA::casA* complementation clones were functionally tested for their ability to grow in salicin, arbutin, and cellobiose. Notably, most complemented Δ*casA::casA* clones matched the growth profiles of their respective WT strains, indicating the full complementation of *casA* function in the cell. However, we also observed that three out of five *K. grimontii* strains, 1/5 *K. oxytoca* and 2/5 *K. michiganensis* strains with PCR-confirmed complementation displayed growth delays or showed no growth on salicin and arbutin (Figure 4a). Overall, our method allows both genetic and phenotypic complementation but underscores that functional validation of complemented strains is critical for these types of genetic studies. Furthermore, *in vitro* growth assays of deletion and complementation strains confirmed the functional homology of *casA* across KoSCs in β-glucoside utilization.

**Figure 4:**
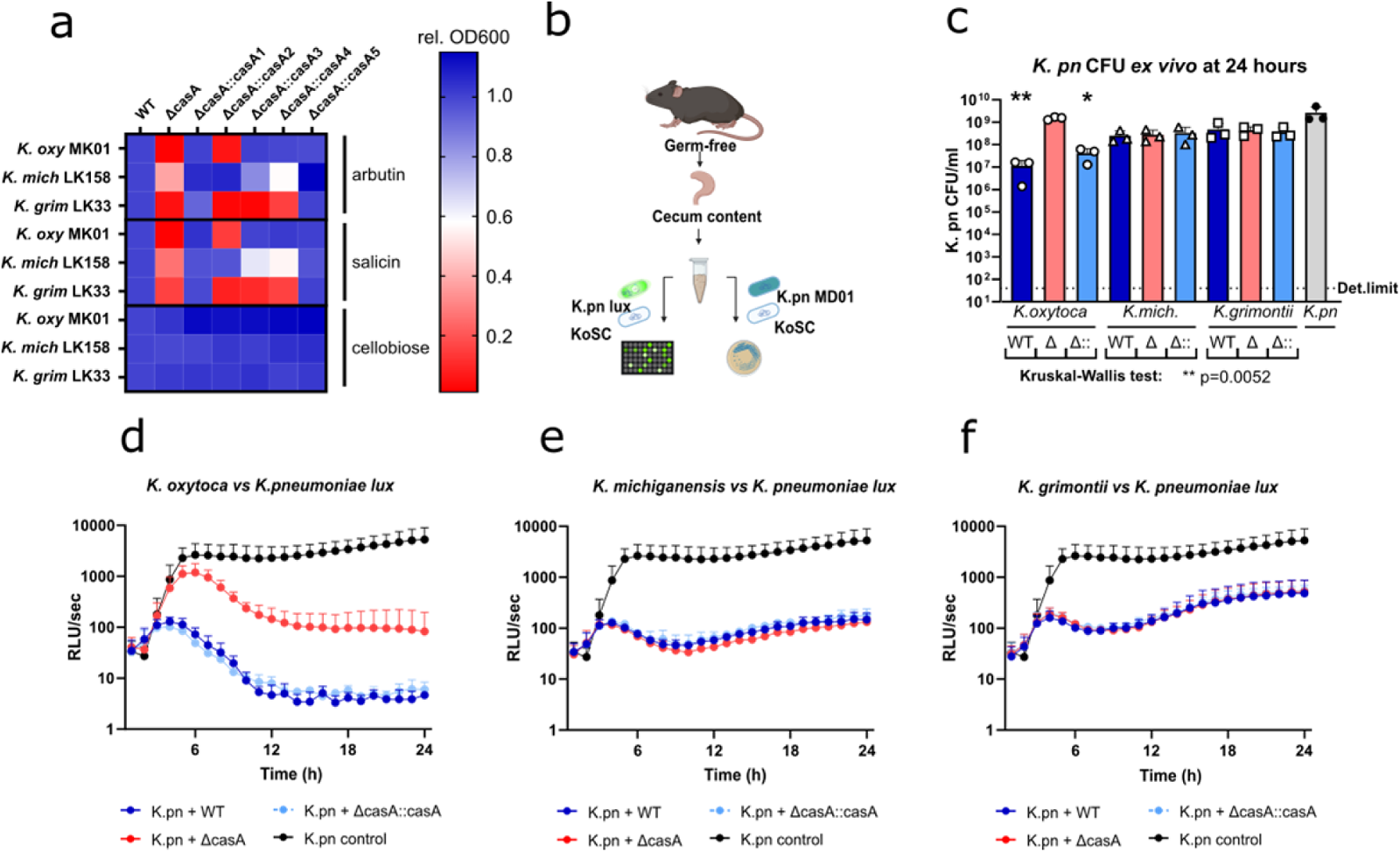
*casA* contributes to *K. pneumoniae* reduction in *K. oxytoca* but not in *K. michiganensis* or *K. grimontii*. **(a)** Heatmap displaying growth on minimal media supplemented with 5 g/L arbutin or salicin. Color scale shows OD_600_ values relative to the WT growth in each condition after 24 hours of incubation. Displayed data represents means of three technical replicates. **(b)** Schematic of *ex vivo* assay using cecum content of germ-free mice in 1:1 dilution with PBS. For longitudinal bioluminescence measurements KoSC members were co-inoculated with *K. pneumoniae lux* carrying the *luxCDABE* operon in the chromosome. Bioluminescence was recorded every hour for 24 hours. For CFU quantifications KoSC members were co-inoculated with MDR human isolate *K. pneumoniae* MR01. After 24 hours of incubation, samples were serially diluted and plated, CFUs were quantified after overnight incubation. **(c)** *K. pneumoniae* MR01 CFUs quantified on LB-agar plates selecting for *K. pneumoniae* growth. Bars represent means of three biological replicates. Each dot represents mean of three technical replicates. p value indicated represents Kruskal-Wallis test on the effect of KoSC presence on *K. pneumoniae* CFUs. p values for each group represents Dunn’s multiple comparision tests compared to *K. pneumoniae* control group with **p < 0.01 and *p < 0.05. **(d-f)** Strength of the bioluminescent signal of *K. pneumoniae* lux longitudinally in the presence of the three KoSC members or alone as a control. Each dot represents means of three independent experiments in technical triplicates using different batches of cecum content obtained from different GF animals for each independent experimen

Finally, to determine whether the presence of *casA* is sufficient for interspecies competition against *K. pneumoniae* across the KoSC, as has been reported for *K. oxytoca*^6^, we utilized an *ex vivo* competition assay using the cecal content of germ-free mice as a media base, as we previously described to reassemble the environment of antibiotic-disturbed gut^6^ (Figure 4b). After 24 h of incubation of KoSC strains with the MDR human isolate *K. pneumoniae* MD01^6^, we quantified *K. pneumoniae* CFUs on selective agar plates. As previously reported, we observed a 100-fold, *casA*-dependent CFU reduction of *K. pneumoniae* in co-culture with *K. oxytoca* MK01, whereas we recorded a moderate 10-fold reduction when co-cultured with *K. michiganensis* and *K. grimontii* (Figure 4c). For time-resolved competition, KoSC strains were co-cultured with a *K. pneumoniae* strain carrying the *luxCDABE* operon^31^ allowing longitudinal recording of the bioluminescent signal, which was used as a proxy for growth. After 12 h, we recorded a 1000-fold reduction in *K. pneumoniae* bioluminescence in co-culture with *K. oxytoca* WT and Δ*casA::casA* strain, which was *casA* dependent, as a lower 50-fold reduction was observed when co-cultured with *K. oxytoca* Δ*casA* (Figure 4d). In contrast, *K. michiganensis* (Figure 4e) and *K. grimontii* (Figure 4f) showed a less pronounced reduction effect of 10-100-fold that was *casA* independent (Figure 4e-f). These results support previous reports on the role of *casA* in growth competition of *K. pneumoniae* with *K. oxytoca* and suggest that this effect is not a general KoSC feature but rather limited to *K. oxytoca*, further highlighting the relevance of *K. oxytoca* in mediating colonization resistance to MDR *K. pneumoniae* species.

## Discussion

Although massive improvements have been made in the development of CRISPR-based and other gene editing technologies, the implementation of such tools is often limited to laboratory strains or model organisms, which can vastly differ from less characterized species or even wild isolates of model organisms in terms of transformability and cultivability^32^. In this study, we present an optimized Cas9-based pipeline for full-length markerless gene deletion combined with a gene knock-in protocol for complementation, enabling downstream functional analyses that do not require the presence of selection markers for plasmid maintenance, which can limit experimental design.

We optimized our pipeline for *K. oxytoca* MK01, which was previously used to delete *casA*^6^. We observed high transformation efficiencies in *K. oxytoca* MK01, making it an ideal candidate for benchmarking our Cas9-based genetic toolbox. Transformation of *K. oxytoca* with sgRNA-targeted Cas9 without induction of the lambda Red system revealed that a small fraction of cells was able to survive or escape the lethal effects of the Cas9-generated double-stranded DNA break, as previously observed in *E. coli*^33,34^ and *P. putida*^19^. These escapers may occur as a result of several possible genetic events interfering with effective gene targeting, such as deletion of sgRNA^19^, mutations in Cas9^33^, mutation of the target site, or homologous recombination-mediated repair in case of weakly targeting sgRNAs^34^. In our study, the number of escapers varied depending on the target sgRNA sequence, although we generally observed strong CFU reductions with all tested sgRNAs.

Comparison of several sgRNAs targeting *casA* (or adjacent genes for *casA* complementation) revealed vast differences in editing efficiencies ranging from 0-100%. While *casA* mutants were obtained for 4/5 sgRNAs, the differences in editing efficiencies highlight the need for testing different sgRNAs in the gene of interest to successfully obtain deletion mutants as well as complementation strains. Notably, in most publications reporting similar CRISPR-based tools, differences between sgRNA performances have rarely been reported^19,21,35^. Furthermore, we observed that sgRNA, which gave rise to more escapers, also allowed better editing when the lambda Red genes were induced, resulting in a higher editing efficiency. This was recently demonstrated in a study that obtained an increased editing efficiency by attenuating sgRNA^29^. While designing multiple sgRNAs can circumvent this heterogeneity for the purpose of gene deletion, in the case of other edits aimed at a specific genomic location, attenuation could potentially be useful for sgRNAs yielding no colonies, such as sg189 (Figure 1d). Another key factor contributing to successful gene editing was the 10-fold increase in L-arabinose concentration for the induction of lambda Red gene expression compared to recently published CRISPR/Cas9 tools in *K. pneumoniae*^21,36^, resulting in an increase in editing efficiencies from 0% to 75% in three previously unsuccessful sgRNAs. This approach is particularly beneficial for hypermucoid strains, which have low transformation efficiency and are common in the KoSC and *Klebsiella* genera in general^37,38^. The increase in the expression of the recombination machinery would circumvent the low transformability and allow the isolation of the desired genetic edit, requiring fewer transformants.

Establishment of robust and high-efficiency editing combined with the implementation of the recently reported ‘bookmark’ complementation method^28^ enabled the development of successful *in situ* complementation, which to our knowledge is reported for the first time for KoSC. Although complementation follows the same principle as deletion, it also poses the challenge of exclusively targeting the chromosome, but not the repair template by Cas9. To circumvent this issue, we first mutated PAM in the repair template, which can be both laborious and inefficient. Adapting a recently reported protocol developed in *Clostridium autoethanogenum*^28^ allowed us to simplify the design process and increase complementation efficiency with unique sgRNAs inserted into the chromosome during gene deletion. Importantly, this led to a similar editing efficiency for complementation as for deletion. After selecting the optimal deletion and complementation conditions for *K. oxytoca* MK01, we successfully applied our approach to four out of five other types and human isolate strains belonging to two additional species in the KoSC, namely *K. michiganensis* and *K. grimontii*.

Finally, the generation of Δ*casA* and Δ*casA::casA* strains in a human-derived strain of each KoSC representative species revealed that *casA*-dependent reduction of *K. pneumoniae* can only be observed in the case of *K. oxytoca*. This demonstrates that the presence of CasA is not sufficient but rather contributes in a *K. oxytoca*-specific manner against colonization by *K. pneumoniae*. Although we observed a moderate, 10-100-fold reduction of *K. pneumoniae* in the presence of *K. michiganensis* and *K. grimontii*, these interactions can most probably be attributed to genes other than *casA*. Hence, more extensive characterization of nutrient competition between members of the KoSC and other Enterobacteriaceae, including *K. pneumoniae*, is required to identify species-versus strain-specific effects.

In summary, we presented a robust and easily applicable two-plasmid CRISPR-Cas9 system for KoSC, enabling full-length gene deletion and subsequent *in situ* complementation. Following gene editing, both plasmids could be easily cured, creating markerless strains for downstream analyses. Given the increasing interest in characterizing competitive interactions between gut bacteria and *Klebsiella* species in particular^5,6,13,39,40^, we anticipate that our tool will contribute to mechanistic studies of microbiome communities.

## Materials and Methods

### Plasmid construction

The single-guide RNA-expressing plasmids were assembled using standard cloning procedures. Briefly, spacers were ordered as oligos with 5′ overhangs corresponding to the BbsI digestion site in the plasmid pSG-GFP-spe. The forward and reverse spacer oligos (10 µM) were heated to 95 °C in a thermocycler, then cooled down gradually at -0.1 °C/s, and finally diluted 1:100. One microliter of annealed oligos was ligated into BbsI-digested pSG-spe following the manufacturer’s instructions, transformed into competent *E. coli* DH5α cells, and plated on LB agar plates supplemented with 75 µg/mL spectinomycin. Following overnight incubation at 37 °C, successfully annealed plasmid bearing colonies appeared white, in contrast to GFP-expressing green colonies.

### Gene targeting in KoSC members

KoSC members were cultured aerobically overnight in LB medium, then back-diluted 50-fold and subsequently grown until cultures reached an OD_600_ of 0.6-0.8. Cells were then made electrocompetent by three rounds of washing with 10% ice-cold glycerol solution and centrifuging at 7.200 g. Competent cells (50 µL) were transformed with 50 ng pCas-apm, and after 60 min of incubation, the cells were recovered in 950 µL liquid LB. The cultures were then centrifuged for 3 minutes at 8.000 g and the bacterial pellets were resuspended in 50 µL liquid LB and plated on LB agar plates supplemented with 60 µg/mL apramycin.

In the second round of transformation, transformants were cultured in liquid LB supplemented with 60 µg/mL apramycin overnight, then back-diluted 50-fold and grown at 30 °C until OD_600_ of 0.15-0.2, then induced with 10% (v/v) 20 g/L L-arabinose in liquid LB supplemented with 60 µg/mL apramycin. Induction continued for 2 h either at room temperature or at 30 °C, depending on the strain’s growth rate – until cultures reached OD_600_ of ca. 0.8. The cells were made electrocompetent as described above.

For genes targeting 50 µL of L-arabinose, competent cells were co-transformed with 200 ng of sgRNA plasmids and ~ 400 ng of linear repair template assembled with 500 bp upstream and downstream homology arms via SOE-PCR. Notably, for bookmark spacer insertion, spacer sequences were included at the 3′ end of primers used for homology arm amplification during oligo synthesis. The cells were recovered after 60 min and plated in serial dilutions on LB agar plates supplemented with 60 µg/mL apramycin and 300 µg/mL spectinomycin. The limit of detection was determined based on the plated volume. Following overnight incubation, colony PCRs were performed on 10 and three randomly selected colonies with targeting and non-targeting sgRNA expressing pSGs, respectively. Successful repair template integration was verified on a 1% TAE agarose gel run at 130 V for 15 min, and then editing efficiencies were quantified.

For complementation and downstream functional analyses, the edited strains were cured with pSG-spe and both plasmids. Briefly, colonies were inoculated under non-selective conditions and then plated on LB agar plates supplemented with 5% sucrose, and incubated at 30 °C or 37 °C depending on pCas-apr curing. Following overnight incubation, the colonies were streaked on selective and non-selective agar plates to confirm successful plasmid curing.

### Bookmark complementation spacer selection

sgRNAs for bookmark complementation must fulfill two important criteria: (1) they must not show targeting activity in the WT genome, which would lead to off-target effects during complementation, and (2) they are suitable for successful gene editing. To eliminate the possibility of off-target activity, we utilized spacer sequences that showed sequence homology to *casA* but contained 2-7 mismatches in all tested KoSC strains along the 20 nt spacer sequence. Furthermore, we compared the targeting activity of all bookmark spacers in uninduced WT KoSC cells to that of a non-targeting spacer.

### Phylogenetic tree construction

The phylogenomic tree in Figure 3a representing the evolutionary relationships in the *Klebsiella oxytoca* species complex was constructed based on the core genes in the indicated seven genomes (242 genes) using FastTree.

### *In vitro* growth assay

To assess the sugar utilization of WT, Δ*casA* and Δ*casA*::*casA* strains of KoSC members, strains were cultured overnight in liquid LB, and then 50 µL of the culture was streaked on R2A agar plates. After 16 h, the cells were adjusted to an OD_600_ of 0.2 in PBS. Cultures were inoculated at 5% v/v in MM9 medium with a single carbon source of arbutin, salicin, or cellobiose at a concentration of 5 g/L. OD_600_ was recorded every hour. Growth assay data were visualized using GraphPad Prism software (version 8.2.1).

### Mice

Germ-free animals were used for intestinal content harvesting in *ex vivo* competition assays. WT C57BL/6NTac mice were raised in a germ-free breeding facility at the Helmholtz Centre for Infection Research and maintained for breeding purposes only. In parallel to renewal of breeding animals, old breeding pairs were sacrificed, and cecum content was isolated, then stored at -20°C until further use.

### Ethics statement

All animal experiments were performed in accordance with the guidelines of the Helmholtz Center for Infection Research, Braunschweig, Germany, the National Animal Protection Law (Tierschutzgesetz, TierSchG), and Animal Experiment Regulations (Tierschutz Versuchstierverordnung, TierSchVersV), and the recommendations of the Federation of European Laboratory Animal Science Association (FELASA).

### *Ex vivo* competition assay

The isolated cecum contents were thawed and diluted 1:1 with PBS. KoSC and *K. pneumoniae* lux strains were cultured in liquid LB overnight, and then the OD was adjusted to OD_600_ of 1 and OD_600_ of 0.2, respectively. 20 µl of KoSC members and 10 µl of *K. pneumoniae* lux was co-inoculated into 250 µL of PBS-diluted germ-free cecum contents in a 96-well format. The bioluminescence was recorded every hour using a microplate spectrophotometer. After 24 h, the assays were plated in serial dilutions on LB agar plates supplemented with 50 µg/mL kanamycin or 50 µg/mL chloramphenicol for *K. pneumoniae* quantification against *K. oxytoca/K. grimontii* and *K. michiganensis* respectively, and on MacConkey agar plates for KoSC quantification of *K. oxytoca* and *K. grimontii*, which are visually distinguishable from *K. pneumoniae* unlike *K. michiganensis*.

## Acknowledgements

We acknowledge Prof. Dr. Christian Riedel (Ulm University) and Prof. Dr. Regis Tournebize (Sorbonne Université, INSERM, U1135) for providing *K. pneumoniae* strains. This work was supported by the Joint Programming Initiative on Antimicrobial Resistance (01KI1824 to C.L.B. and T.S.) and the BMBF (DF-AMR2: 01KI2131 to T.S.). The funders had no influence on the study design, data collection or analysis, or publishing process.

## Data availability statement

The authors confirm that data supporting the findings of this study are available within the article.

